# Generative adversarial networks simulate gene expression and predict perturbations in single cells

**DOI:** 10.1101/262501

**Authors:** Arsham Ghahramani, Fiona M. Watt, Nicholas M. Luscombe

**Affiliations:** The Francis Crick Institute, 1 Midland Road, London NW1 1AT, United Kingdom; King’s College London, Centre for Stem Cells and Regenerative Medicine, 28th Floor, Tower Wing, Guy’s Hospital, Great Maze Pond, London SE1 9RT, United Kingdom; UCL Genetics Institute, University College London, London WC1E 6BT, United Kingdom; Okinawa Institute of Science & Technology Graduate University, Okinawa 904-0495, Japan

## Abstract

Recent advances have enabled gene expression profiling of single cells at lower cost. As more data is produced there is an increasing need to integrate diverse datasets and better analyse underutilised data to gain biological insights. However, analysis of single cell RNA-seq data is challenging due to biological and technical noise which not only varies between laboratories but also between batches. Here for the first time, we apply a new generative deep learning approach called Generative Adversarial Networks (GAN) to biological data. We apply GANs to epidermal, neural and hematopoietic scRNA-seq data spanning different labs and experimental protocols. We show that it is possible to integrate diverse scRNA-seq datasets and in doing so, our generative model is able to simulate realistic scRNA-seq data that covers the full diversity of cell types. In contrast to many machine-learning approaches, we are able to interpret internal parameters in a biologically meaningful manner. Using our generative model we are able to obtain a universal representation of epidermal differentiation and use this to predict the effect of cell state perturbations on gene expression at high time-resolution. We show that our trained neural networks identify biological state-determining genes and through analysis of these networks we can obtain inferred gene regulatory relationships. Finally, we use internal GAN learned features to perform dimensionality reduction. In combination these attributes provide a powerful framework to progress the analysis of scRNA-seq data beyond exploratory analysis of cell clusters and towards integration of multiple datasets regardless of origin.

## 1. Introduction

The development of affordable single cell RNA-seq has enabled the measurement of transcript abundance in hundreds to thousands of individual cells^1–3^. These profiles provide an opportunity to define cell state and indirectly measure the signals and factors influencing cell fate. However, despite the richness of single cell measurements, these data are computationally challenging to analyse due to technical and biological noise meaning that traditional bulk expression analysis approaches are frequently not applicable. Current single cell RNA-seq dimensionality reduction methods can successfully reveal clustering and structure within data when technical noise is low; however, they cannot easily integrate diverse datasets produced using distinct protocols^4–6^. Furthermore, current approaches focus on differential expression and marker gene identification but do not yield functional gene regulatory relationships.

Deep learning algorithms enabled by advances in computational power have demonstrated the capability to analyse diverse datasets from images to genomics^7,8^. In particular, generative adversarial networks (GAN), first introduced in 2014, have shown promise in the field of computer vision^9^. Since their introduction, GANs have become an active area of research with multiple variants developed that have resulted in improved performance and training^10–14^. Common to all of these variants^15^ is the concurrent training of two neural networks competing against one another, referred to as the generator and discriminator (Figure 1). The generator is tasked with generating simulated data, whereas the discriminator is tasked with evaluating whether data is authentic or not. Notably, only the discriminator directly observes real data while the generator improves its simulations through interaction with the discriminator. As training progresses both neural networks learn key features of the training data. Additionally, both discriminator and generator performance improves as they compete against each other.

**Figure 1.**
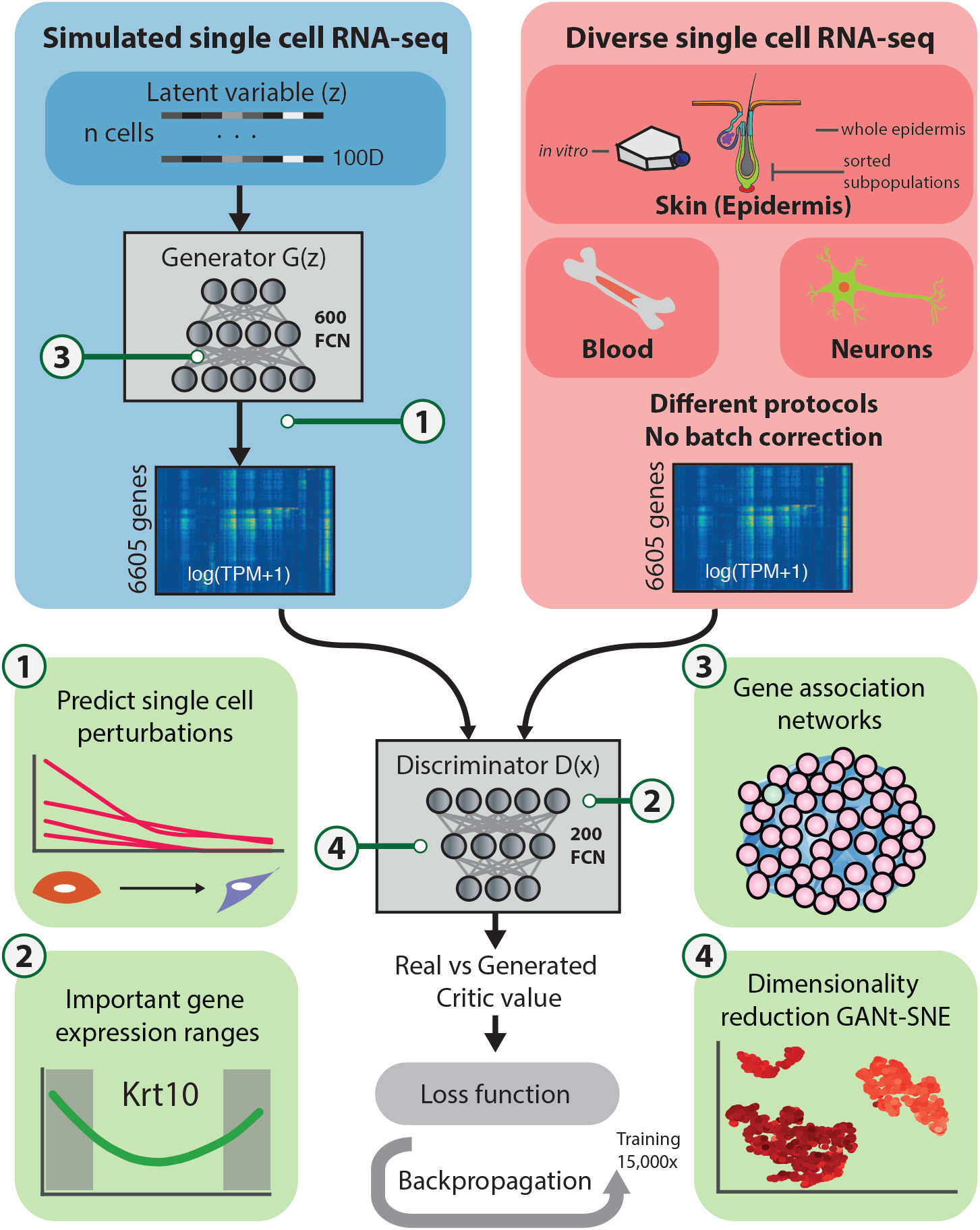
Overview of single cell RNA-seq analysis with generative adversarial networks. Generative adversarial networks consist of two neural networks concurrently training whilst competing against one another. These networks referred to as the generator and discriminator networks each have a distinct task. The generator is tasked with generating data by transforming a 100-dimensional latent variable into a single cell gene expression profile. In turn the discriminator is tasked with evaluating whether data is authentic or generated. Only the discriminator network sees the real single cell RNA-seq data, which have not been corrected for batch effects or technical variation. Once trained, we are able to extract biologically meaningful information from the generator and discriminator networks such as gene association networks, predict single cell time-courses, important gene expression ranges and biologically meaningful dimensionality reduction. Fully connected neurons, FCN.

In the field of computer vision, GANs have proved capable of generating visually convincing and novel images such as faces or furniture^16,17^. Furthermore, it has been shown that the generator network learns a meaningful latent space where visually similar data such as similar faces are clustered. A meaningful latent space for single cell RNA-seq is particularly appealing as this can be used in conjunction with existing dimensionality reduction methods for improved and more meaningful subpopulation extraction. Additionally, an advantageous feature of generative neural networks is that non-linear relationships between features of the data are learned through training and can be later extracted to provide further insights.

Here, we show that GANs are able to integrate datasets produced using distinct scRNA-seq protocols and by separate laboratories. We demonstrate this by applying the GAN to mouse epidermal, mouse neural and human hematopoietic single cell RNA-seq data. These datasets were chosen to span a range of biological and technical variation, including single cells isolated using FACS, the Fluidigm C1 microfluidics platform and the 10x Genomics Chromium platform in separate labs. We focus analysis and exploration on epidermal data and integrate three disparate datasets. The mammalian epidermis comprises interfollicular epidermis (IFE), hair follicles, sebaceous glands and further associated adnexal structures: together they form a protective interface between the body and external environment. Hence, epidermal cell fate is determined through the integration of extracellular cues with transcriptional activity^18,19^. Under steady-state conditions, each epidermal compartment is maintained by distinct stem cell populations. However, under certain conditions such as wounding, each stem cell subpopulation is able to contribute to all differentiated lineages^20^. These distinct yet seemingly interchangeable subpopulations hint at a common, but as yet undefined epidermal gene regulatory network.

For the first time, we have applied GANs to genomic data to uncover previously unknown gene regulatory relationships and regulators of epidermal cell state. Using the trained generator neural network we demonstrate that GANs can be used to predict the effect of cell state perturbations on unseen single cells. We further show that GANs produce biologically meaningful dimensionality reduction. Although existing algorithms are able to perform these analyses in isolation, we demonstrate that, once trained, GANs are able to learn all relevant features for these analyses whilst concurrently providing a generative model of scRNA-seq.

## 2. Results

### 2.1 Generative adversarial networks integrate diverse datasets

We first used a generative adversarial network (GAN) to integrate multiple diverse mouse epidermal single cell RNA-seq datasets. These datasets originated from three labs, spanning mouse epidermal cells *in vitro^21^*, whole epidermis *in vivo^22^* and isolated subpopulations *in vivo^23^* (Figure 1, upper right. See methods for GEO accession numbers). No adjustments were made to account for batch-to-batch variations within datasets and across labs. After removing non-epidermal cell types and outlier cells we retained 1,763 cells for training and 500 unseen cells for testing and validation. Similarly to other analysis methods, we filtered genes by expression level, resulting in 6605 potentially informative genes (see methods).

GANs consist of two neural networks competing against each other. The generator is tasked with producing realistic output data from a random input vector z or latent variable. In our case the latent variable z is of lower dimension (100, arbitrarily chosen) than the output scRNA-seq data (6605 genes), hence the generator represents a mapping from a lower dimensional space to gene expression space. Each 100-dimensional latent vector maps to 6,605 transcripts per million (TPM, log(TPM+1)) normalised gene expression values for one cell. The second network, named the discriminator, is tasked with discriminating between the real and generated data. The discriminator takes log(TPM+1) gene expression values as input and outputs a score related to its assessment of the gene expression input. These two networks are trained using a combined loss function leading to improved performance of both neural networks (Figure 1).

We tested several variations of GANs ultimately utilising a Wasserstein GAN with added gradient penalty term in the loss function as described previously^24^ due to training stability. We trained our neural network for approximately 15,000 epochs per full training run. We evaluated the generator network output performance at multiple checkpoints using t-distributed stochastic neighbour embedding (t-SNE, Figure 2A) and correlation between real samples and generated samples as shown in Supplementary Figure 1. For each evaluation we generated 500 cells using 500 random latent variables. Early in training, the GAN struggles to produce a varied output representative of the breadth of cell types and states; generated cells are closely correlated and form a single cluster in the t-SNE plot (Figure 2A 1000 steps, Supplementary Figure S1A). After 5,000 steps the generator begins to produce a varied output with generated cells mapping to multiple clusters in the t-SNE plot covering the different cell types, cell states, cell origin and experimental batches present in our combined dataset. After 10,000 steps, the generator network is capable of producing a subset of cells with similar transcriptomic profiles to those of Yang and colleagues that form a distinct cluster. Further GAN training broadens the distribution of correlations between generated samples, indicating an increasingly diverse generator output (Supplementary Figure S1A). We observed that after 13,000 steps continued training does not continue to increase the output diversity of the generator network, as defined by the median distance between cell gene expression profiles. Figure 2B shows that GAN output diversity reaches a maximum between 12,500 and 15,000 training steps. Furthermore, additional training between 12,500 and 15,000 steps does not change the generated cell correlation distribution (Supplementary Figure S1A). At this point the discriminator loss function has also converged and therefore we cease further training (Supplementary Figure S2).

**Figure 2.**
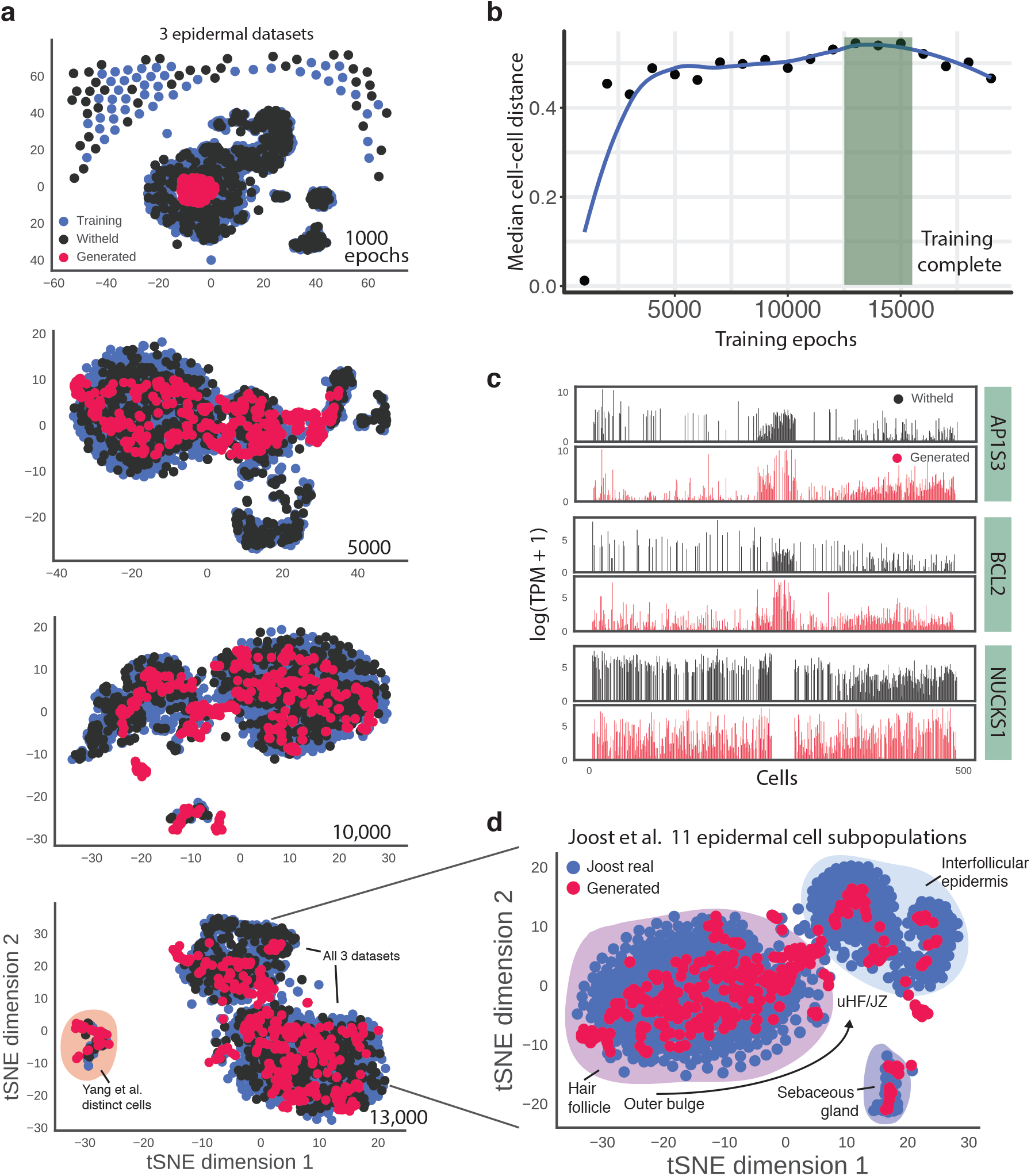
GANs model epidermal single cell gene expression. **a**) t-SNE projection of 1763 real training cells, 500 real withheld cells and 500 generated single cell expression data: each data point represents a cell and cells with similar expression profiles are positioned close together. Generated cells are clustered at the beginning, but with training gradually occupy the entire range of expression outputs for different cell types across three studies. Cells are visualised at 1000, 5000, 10,000 and 13,000 training steps. **b**) Relationship between GAN training epochs and output diversity as measured by median cellcell distance. Blue line is a LOESS regression. The greatest diversity of gene expression is achieved between 13,000 and 15,000 steps. **c**) Bar plots showing expression of Ap1s3, Bcl2 and Nucks1 in 500 unseen real cells and 500 closely related generated cells. Cells are ordered by hierarchical clustering of the unseen real cells using all 6605 genes. **d**) t-SNE projection of subset of real cells from Joost et al. and corresponding generated cells. Despite training with a larger and diverse dataset, the GAN captures cell diversity at a detailed level described by Joost. Labels show the corresponding cell type for each cluster. Upper hair follicle, uHF. Junctional zone, JZ

There is currently no standard method for assessing generative models of gene expression data. We computationally validated our GAN by examining the correlation between generated samples and unseen samples (Supplementary Figure S1B). The distributions obtained at different training steps show that generated samples are correlated with, but distinct from, real samples. At early training steps this distribution is centred around a Pearson correlation of 0.4. After 10,000 steps this distribution broadens, indicating the generator learns to simulate a diverse population of cells. This can also be seen from the expression of individual genes. Figure 2C shows single cell expression bar plots for three genes that distinguish subpopulations in our cell cohort, Ap1s3, Bcl2 and Nucks1. For each gene we show 500 unseen cells and 500 closely related generated cells with an average correlation of 0.71. Our generated cohort follows a similar pattern of expression to the real unseen cells, with Ap1s3 and Bcl2 highly expressed in differentiating cell types, and Nucks1 expression absent in vitro. In real cells where Ap1s3 or Bcl2 are not detected, the corresponding generated cells show low non-zero expression indicative of a form of imputation performed by the generator. Across all genes, expression variability in the simulated cells is similar to the real data. Together these results demonstrate that the generator network is not memorising and reproducing training samples but is instead inferring relationships between gene expression values in order to output convincing heterogeneous generated cells.

We noted that generated subpopulations of cells overlap with real populations, however, for most subpopulations the generated cells do not cover the entire variation within a subpopulation (see upper cluster in Figure 2A, 13,000 epochs). One contributing factor is the discrete drop-out nature of the real single-cell data in comparison to generated cells. Real scRNA-seq expression values are derived from discrete read count data whereas the generator network consists of a continuous input (latent space, *z)* and resulting continuous output which approximates the real discrete gene expression distribution. This property of the generator network results in overlapping but less variable subpopulations of cells.

Focusing on the data from Joost and colleagues, the 1422 cells (one of the three integrated datasets) comprise eleven epidermal cell types originating from whole dorsal epidermis including previously uncharacterised cells. The generator neural network is able to simulate cells clustering with all *in vivo* epidermal cell types, ranging from sebaceous gland cells with distinct gene expression profiles to upper hair follicle and interfollicular epidermal cells (IFE) with similar transcriptomes but distinct spatial positions (Figure 2D). This range of cell type simulation is achieved when examining generated cells at multiple resolutions despite training the GAN using three diverse datasets.

### 2.2 Generative modeling of large and sparse scRNA-seq datasets

We next applied the GAN algorithm to non-epidermal datasets to validate the methods ability to generalise. We focused on the generative capabilities of the GAN and assessed whether it was equally capable of generating cells from non-epidermal datasets. We selected three datasets on the basis of their diversity.

First, we applied the GAN to a timecourse of motor neuron differentiation^25^. Briefly, Sagner and colleagues started from mouse embryonic stem cells (ESCs) and over seven days directed their differentiation towards motor neurons using a combination of signalling factors including Wnt and FGF. Single cells were isolated and libraries prepared using the Fluidigm C1 platform. For this dataset we trained the GAN using 2,343 cells and 14,000 genes, substantially more than for the epidermal datasets owing to the more diverse range of cell types present from ESC to motor neuron. Figure 3A shows a t-SNE projection of real and generated cells resulting from training the GAN on this motor neuron dataset. The GAN is able to generate cells from all cell type clusters present in the real dataset and produces a similar number of outlier cells to the real scRNA-seq.

**Figure 3.**
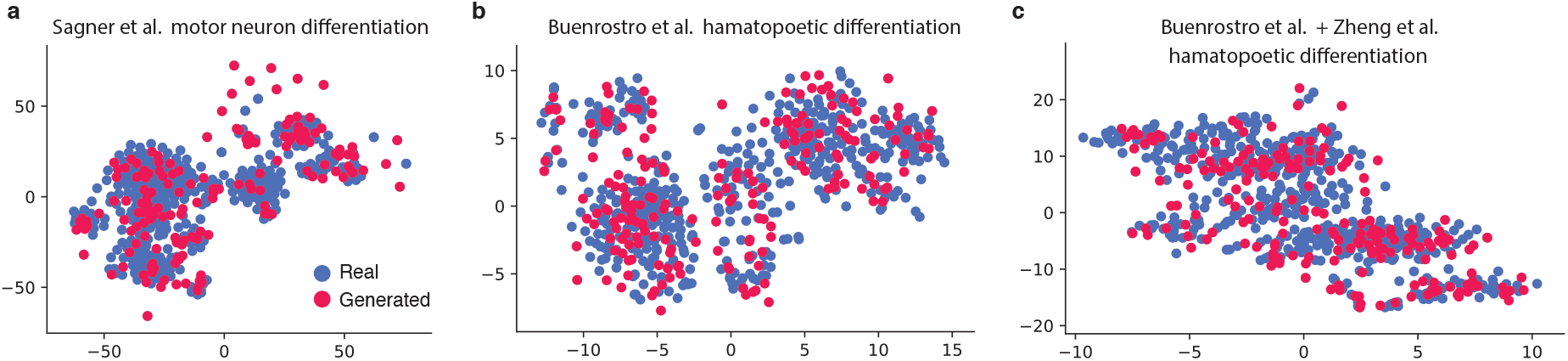
Generative modeling of diverse scRNA-seq datasets. **a-c**) t-SNE projection of 500 real cells and 300 GAN-generated cells after training on scRNA-seq profiles of mouse neural progenitors from Sagner et al. (a), human hematopoetic cells from Buenrostro et al. (b), and a combined dataset of human hematopoetic cells and FACS-sorted monocytes from Zheng et al. (c)

Second, we applied the GAN to two droplet-sequencing based single cell datasets. We first analysed 7,108 cells from Greenleaf and colleagues alone^26^. This dataset consists of CD34+ cells isolated by FACS from human bone marrow which includes cell types spanning the myeloid, erythroid, and lymphoid lineage. Subpopulations cover the full spectrum of human hematopoietic differentiation comprising a minimum of 10 phenotypically distinct cell types sequenced using the 10x Genomics Chromium system. A key feature of the Chromium system is the high-throughput of sequenced cells, enabling profiling tens of thousands of cells. Furthermore, sequenced libraries are of lower depth than FACS or Fluidigm C1 based methods with typically 50-100,000 aligned reads per cell in contrast to up to 1 million for the Fluidigm C1 epidermal data from the Joost dataset. A median of 4,400 genes were detected per cell. As before, the fully trained generator neural network is able to generate cells clustering with the full spectrum of gene expression profiles. Increased sparsity of the data and reduced number of detected genes per cell did not appear to affect GAN training. We next retrained on the combined dataset of Buenrostro and Zheng, a publicly available dataset profiling CD34+ and CD14+ monocyte cells^27^ (13,034 cells in total) to understand whether the trained generator network could successfully simulate gene expression profiles from two large datasets. Figure 3C shows the t-SNE projection after GAN training on the combined dataset. Here again, one trained generator network demonstrates the ability to simulate cells clustering with two datasets and two distinct sets of technical and biological variation.

### 2.3 Simulating cellular perturbations using latent space interpolation

To take advantage of the gene expression rules learned by the generator network we devised an algorithm for retrieving the latent space vector *z* from an arbitrary gene expression profile x such that *x* = *G*(*z*). For each target cell we randomly sampled latent space vectors (*z*) and generated cells until a sufficiently similar cell is obtained (see methods). In other fields where GANs have been applied, such as image generation, the latent space representation of an image meaningfully represents visually similar images. Furthermore, vector arithmetic in the latent space leads to meaningful outputs. For example Radford and colleagues^10^ have shown that subtracting the latent space vectors of a face wearing glasses from a face without glasses results in a differential vector representing glasses; adding this to a different face outputs a face with glasses (i.e.G(*z*′) produces a person wearing glasses where *z*′ = (*z_glasses_ - z_no gasses_*) + *z_face_*).

We hypothesised that latent space arithmetic can be extended to cell types and cell states. To investigate this we simulated the process of epidermal differentiation at the single cell level using latent space arithmetic, as shown in Figure 4A. We sampled 30 terminally differentiated and undifferentiated pairs of cells from the unseen pool of cells in the Joost dataset using the original study labels, obtained their latent space vectors *z_differentiated_* and *z_basal_* and calculated the difference between these vectors *δ* = *z_differentiated_ - z_basal_*. We sampled cells from both the interfollicular epidermis (IFE) and hair follicles (HF) in order to obtain a universal differentiation latent vector. Sampling from both IFE and HF subpopulations allows us to construct a latent space representation of differentiation capturing the common differentiation-associated gene expression changes and concurrently averaging out location-specific effects. *In vivo* both compartments are able to repopulate each other under wound healing conditions and undergo similar cellular changes during differentiation. Hence, it is reasonable to assume there is a common gene regulatory programme driving HF and IFE differentiation which can be captured by a common latent space vector. Unseen cells were used to demonstrate that the generator’s learned latent space could generalise to previously unseen gene expression profiles. We then added *δ*, the latent space differentiation vector, to an unseen undifferentiated cell and interpolated 1000 timepoints between the undifferentiated latent space point and the simulated differentiation latent space endpoint. We used the generator network to produce gene expression for all 1000 timepoints and repeated this process for 30 IFE cells with similar undifferentiated starting profiles. It is important to note that only the starting point (the undifferentiated cell gene expression profile) is real, whereas the differentiated endpoint and all timepoints in between are generated by the neural network.

**Figure 4.**
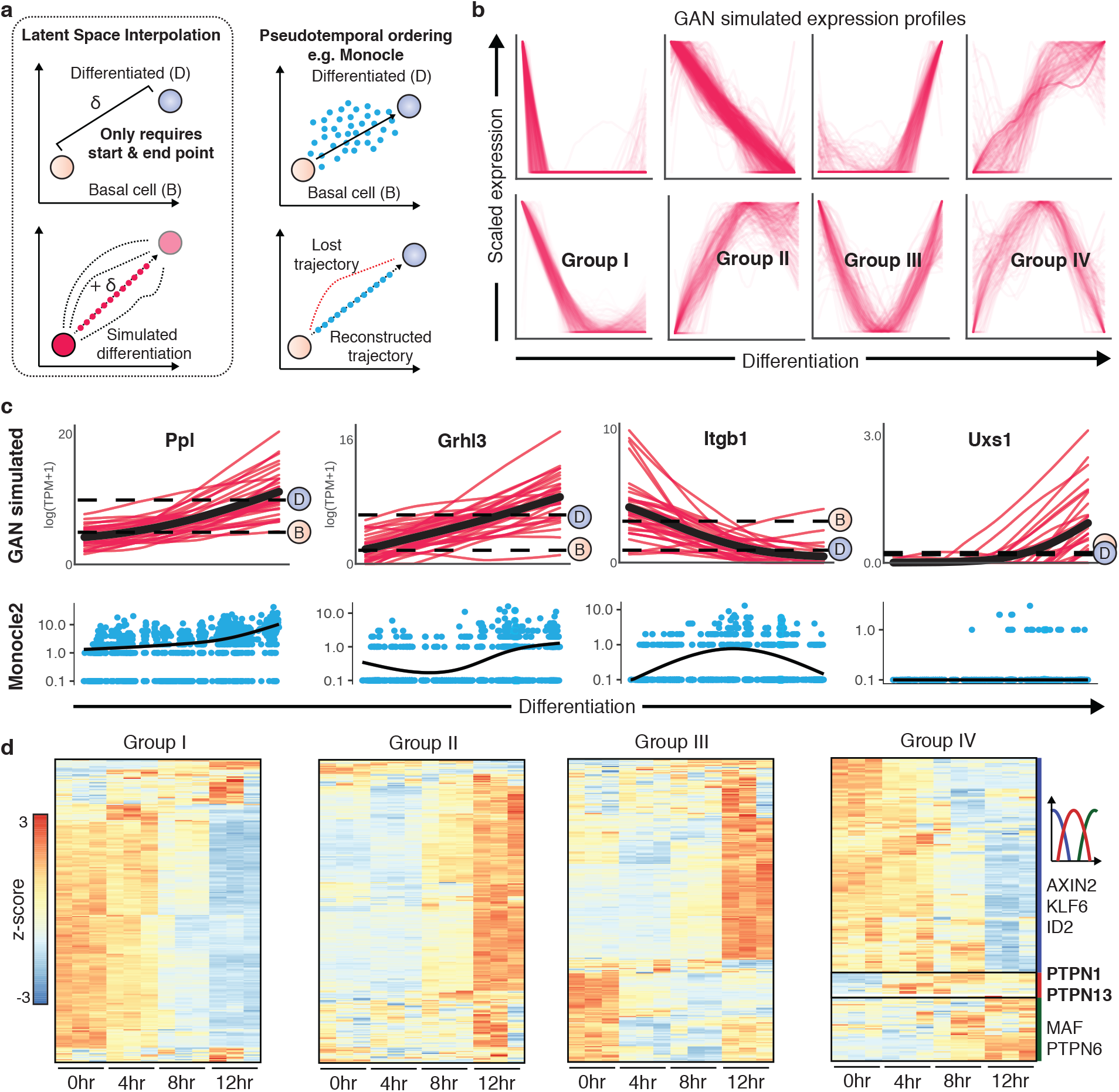
Latent space interpolation allows simulation of single cell differentiation. **a**) Schematics showing how to simulate differentiation for single cells using latent space interpolation. Expression data for pairs of undifferentiated and differentiated cells are transformed to their respective latent space vectors (*z_basal_, z_differentiated_*) The difference between the vectors (*δ*) represents differentiation in latent space (upper panel). This difference vector is then applied to the latent space vector of another unseen cell in order to predict the terminally differentiated state (lower panel). We also simulate 1000 differentiation timepoints. **b**) K-means clustering of simulated differentiation expression profiles for 6605 genes. We identify eight broad types of temporal gene expression, of which four of the dynamic expression patterns are labelled Groups I-IV. **c**) GAN-simulated expression profiles for four representative genes in 30 cells over 1000 differentiation timepoints (upper row, red). Red lines indicate expression profiles in individual cells as predicted by the generator network and black lines provide the mean expression over all simulated cells. Dotted lines correspond to the median expression of basal undifferentiated (B) and differentiated (D) cells from the Joost dataset. Lower row shows expression along differentiation as predicted by Monocle pseudo-ordering where blue points are individual cells and the black line shows the reconstructed mean expression level. Expression dynamics are shown for two known differentiation markers: Periplakin (Ppl) and Grainyhead like transcription factor 3 (Grhl3), an IFE stem cell marker, Integrin beta-1 (Itgb1) and UDP-Glucuronate Decarboxylase 1 (Uxs1), a gene with unknown epidermal function which displays non-linear expression profiles with substantial cell to cell variability. See Supplementary Figure S5 for further examples. **d**) Heatmaps showing bulk expression levels of genes in Groups I-IV from Mishra et al. at 0, 4, 8 and 12 hours of suspension-induced differentiation. Most genes display similar expression profiles to the simulations, despite the differences in experimental set ups.

To investigate the dynamics of gene expression over epidermal differentiation we clustered simulated time-series profiles for all 6605 genes. Using k-means clustering we were able to distinguish eight types of dynamic expression profile (Figure 4B). Genes expressed early in differentiation were enriched for epidermal stem cell related gene ontology and concordantly late-expressed genes were enriched for epidermal differentiation (q-value<0.01 for both, Supplementary Figure S3).

Figure 4C shows simulated time-series gene expression profiles for a selection of genes. Two differentiation markers, Periplakin (Ppl) and Grainyhead-like protein 3 homolog (Grhl3), demonstrate the ability of our latent space arithmetic to simulate successfully simulate differentiation of single cells. Over differentiation time points the mean expression level of these two markers increases, spanning the expression levels observed in the undifferentiated and differentiated subpopulations (B - basal start point, D - differentiated end point median expression line). Similar to real scRNA-seq gene expression profiles, the generated time points for individual cells display substantial cell-to-cell variation in gene expression over time, with cells showing different gene expression curves dependent on initial expression level and cell state. Similarly, for basal IFE marker such as Integrin-beta 1 (Itgb1, Figure 4C) and Metallothionein 2A (Mt2, Supplementary Figure S5) the reduction in expression is successfully captured by the GAN latent space interpolation.

Moreover, the expression curve is not necessarily monotonic or linear. We also observed a r
ange of highly non-linear expression profiles such as genes only expressed in early or late timepoints (Uxs1), parabola-like expression (Actb, Ube2f, Supplementary Figure S7) and genes which change in expression variation but not mean expression. For the example of Uxs1, expression is only observed in a small number of cells late in differentiation. A pseudo-ordering or averaging approach would not predict differentiation-induced Uxs1 expression as Uxs1 + cells would be outnumbered by the majority of Uxs1^−^ cells. These generated non-linear expression profiles demonstrate that latent space interpolation produces meaningful gene expression predictions based on gene expression interdependencies and does not simply average between gene expression profiles.

Next, we applied Monocle to the Joost dataset to contrast our predicted gene expression profiles to pseudo-ordering derived observations. Monocle performs a dimensionality reduction of cells and orders gene expression profiles by similarity in the dimensionally reduced space in order to obtain a pseudo-time reconstruction of gene expression over time. Pseudo-ordering produced one single average gene expression trajectory, in contrast to our GAN simulation which produces many equally likely trajectories. We applied Monocle to IFE cells only in order to capture IFE differentiation dynamics and used the in-built Census algorithm for mRNA abundance quantification. Supplementary Figure 5 shows the predicted cell state trajectories when Monocle is applied using default settings. Using this cell ordering, we reconstructed gene expression dynamics for Ppl, Grhl3, Itgb1 and Uxs1 as shown in Figure 4C. For Ppl and Grhl3 we expect increasing expression over differentiation. Monocle’s pseudo-ordering captures this increase, however, the reconstructed expression dynamics over differentiation are constructed from the mean of ordered cells. Hence, the Monocle method predicts one possible gene expression trajectory over time and is not able to predict the likely difference in dynamics from cell to cell. One advantage of a generative model of gene expression is the ability to simulate gene expression dynamics for many cells.

**Figure 5.**
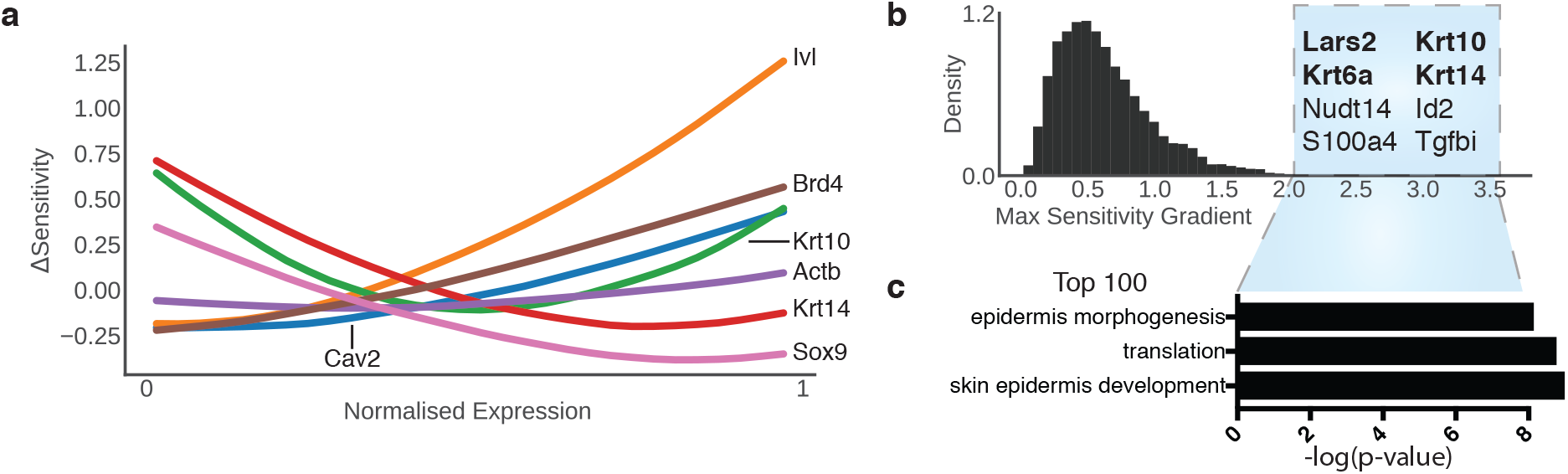
Sensitivity analysis identifies state-determining genes and local gene association networks. **a**) Sensitivity curves for seven known markers for epidermal or determinants of epidermal state: Actin beta (Actb), Involucrin(Ivl), Keratin 10 (Krt10), Keratin 14 (Krt14), Caveolin 2 (Cav2), Bromodomain-containing protein 4 (Brd4), SRY-Box 9 (Sox9). Some genes such as Ivl an IFE differentiation marker display higher sensitivity at increased expression levels whereas others display the opposite trend. **b**) Histogram of maximum sensitivity curve gradients of 6605 genes, with the top eight genes labelled; known markers are highlighted in bold. **c**) Gene ontology enrichment for the top 100 genes by maximum sensitivity gradient as scored by −log10(p-value).

Focusing on Itgb1, we expect expression of this basal integrin to decrease over differentiation. Here, Monocle pseudo-ordering has mis-ordered cells early and mid-differentiation, hence failing to capture that cells earliest in differentiation highly express Itgb1. This can be partially rectified by guiding pseudo-ordering using Itgb1 as a marker gene. Finally, few cells express Uxs1, hence Monocle predicts that Uxs1 is invariant to differentiation, as the Uxs1 + cells are averaged out by the majority of Uxs1^−^. Observing the pseudo-ordering it is apparent that Uxs1 is only expressed by differentiating cells, a feature captured by the GAN latent space interpolation.

To further investigate our gene expression predictions, we focused on four clusters of dynamic gene expression profile, two corresponding to increasing and decreasing expression (Figure 4B, Groups I and II) and two subsets of genes only transiently up- or down-regulated during commitment to differentiation (Figure 4B, Group III and IV genes). To validate these findings we used a recently published dataset investigating human epidermal differentiation^28^. Mishra and colleagues utilised methylcellulose suspension-induced differentiation to obtain temporal differentiation gene expression data from human keratinocytes at 0, 4, 8 and 12 hours. A majority of our predicted transiently expressed genes are dynamically expressed in this dataset with Groups I and II showing decreasing and increasing bulk expression over the differentiation time course (Figure 4D, Group I and II). Focusing on Group IV, these 223 genes are of particular biological interest as they are predicted to be transiently expressed during commitment to differentiation and are therefore likely to play a functional role in this process. From our predicted epidermal commitment genes, this group clusters into three subgroups based on bulk peak expression around 0 hours, between 4 and 8 hours, and 12 hours or later. We hypothesise that three groups are observed as our simulated differentiation assumes a synchronised population of differentiating cells. The asynchronous nature of *in vitro* differentiation and heterogeneous starting population reduces our ability to detect transiently expressed and subsequently downregulated genes. From the subgroup with peak expression between 4 and 8 hours we identified two protein phosphatases, PTPN1 and PTPN13 which are also identified and extensively validated by Mishra and colleagues. This is followed by transient MAF expression - a member of the AP1 subfamily of differentiation transcription factors - predicted by our latent space interpolation approach and observed in the Mishra dataset along with previous studies^29^.

In summary, using our generative model we have simulated a perturbation to cell state in the form of epidermal differentiation and subsequently obtained high-resolution predicted gene expression profiles. Using the generator neural network we have obtained information on switch-like expression of genes, an aspect of gene expression which is difficult to infer from previous methods of single cell RNA-seq analysis such as pseudotime ordering. Pseudo-ordering of cells provides an understanding of the sequence of gene expression events, however, this can be confounded by highly transient gene expression events which may lead to erroneously missing or identifying cell state transitions. In comparison our generative method is guided by gene expression rules learned by the GAN, hence producing valid gene expression profiles at all timepoints. These results can be extended to any other perturbation captured by latent space.

### 2.4 Discriminator network identifies state-determining gene expression ranges

As GAN training progresses, the discriminator network learns to identify compatible ranges of gene expression levels and interrelationships for all genes. An advantage of our approach is that these relationships can be highly non-linear, for example the discriminator can learn gene expression interdependencies which only apply when genes are expressed in a certain range or combination. To extract this learned information we performed a sensitivity analysis on the discriminator network by taking real gene expression profiles from the unseen cell group and varying expression of genes individually from the lowest observed expression level in the cohort to the maximum observed expression level. Figure 5A shows the relationship between adjusted expression level and change in discriminator network output critic value. This analysis produced a sensitivity curve for each gene where absolute sensitivity value indicates a strong change in discriminator critic value and the gradient of the sensitivity curve denotes ranges of gene expression that the discriminator is sensitive to. For genes with no known role in epidermal cell state such as Actb there is little change in sensitivity across all expression levels. In contrast, for known markers or determinators of epidermal state, such as Ivl, Krt10, Krt14, Cav2, Brd4 and Sox9, there is a strong relationship between expression level and sensitivity. For some genes such as Ivl, an IFE differentiation marker, an increase in expression showed increased sensitivity, i.e. a greater effect on discriminator critic value. However, for other genes such as Krt10 medium levels of transcription resulted in a lower sensitivity than low and high levels indicating that for this gene there are two important ranges of transcription. The discriminator network identification of Krt10 import expression ranges is supported by strong expression of Krt10 in the interfollicular epidermis above the basal layer and subsequent downregulation in terminally differentiating cells^22^. Krt10 misexpression has been shown to cause epidermal barrier defects^30,31^. Furthermore, a cellular need for binary expression of Krt10 is clear as the protein constitutes approximately 40% of all cellular proteins in the suprabasal layer of the epidermis^32,33^.

We sought to compare the relative importance of genes in determining cell transcriptional state across all epidermal cell types. Sensitivity varies non-linearly with expression level, hence, to compare all genes we calculated the maximum gradient of the sensitivity curve for each gene (Figure 5B). This metric is an indication there is at least one range of expression levels which has a large effect on discriminator network output. Strikingly, the top 100 genes identified are highly enriched for known epidermal regulators and comprise both low and highly expressed genes (Figure 5C). Amongst the top 10 are three keratin genes with spatially-restricted expression; Krt6a (inner bulge), Krt10 (suprabasal IFE) and Krt14 (basal IFE). Our analysis predicted Lars2 expression as one of the most important indicators of pan-epidermal cell state. Lars2 is a mitochondrial leucyl-tRNA synthetase and is upregulated during differentiation as seen by our differentiation predictions (Supplementary Figure S5) and from the Joost dataset^22^. Considering these analyses we hypothesise that Lars2 is one of the key regulators of cell metabolic considerations during differentiation. This sensitivity analysis approach enables identification of state-regulating genes without bias for transcript abundance.

### 2.5 GAN-derived gene association networks predict Gata6 targets

Finally we examined the internal features of the generator network to extract gene expression interdependencies to complement our sensitivity analysis ranking of state-determining genes by providing context on how these genes are coregulated. The final layer of the generator network non-linearly transforms an arbitrary number of internal features to the final 6605 gene expression values using a leaky rectified linear unit activation function. We found empirically that 600 internal features produced stable and diverse generator output, representing a maximum of 600 features that, when non-linearly combined, produce each gene expression output value. Since these internal features are an order of magnitude fewer than the number of gene expression values, the generator network is forced to learn the most salient relationships between genes in order to produce a convincing output. Hence, genes with correlated generator final layer values are predicted to correspond to co-regulated genes. This is distinct from correlation analysis of gene expression, which infers linear and directly correlated regulation.

We compared GAN-derived gene regulatory associations with expression co-correlation and GRNBoost2, the algorithm employed by SCENIC^34^. GRNBoost2 uses a Random Forest based approach to identify non-linear gene regulatory dependencies from gene expression data. Figure 6A visualises gene-gene association scores derived from these three methods. We observed that gene-gene relationships obtained from the GAN and SCENIC were both more sparse in comparison to co-correlation, with each transcription factor typically strongly associated with fewer than 20 genes. To compare the similarity of identified regulators from these methods, we focused on 512 transcription factors and calculated the overlap between genes in the top 50 predicted targets for each transcription factor (Figure 6B). We found a statistically significant overlap between targets identified from all three methods, indicating that the generator neural network has learnt linear and non-linear gene regulatory relationships similar to GRNBoost2 and co-correlation.

**Figure 6.**
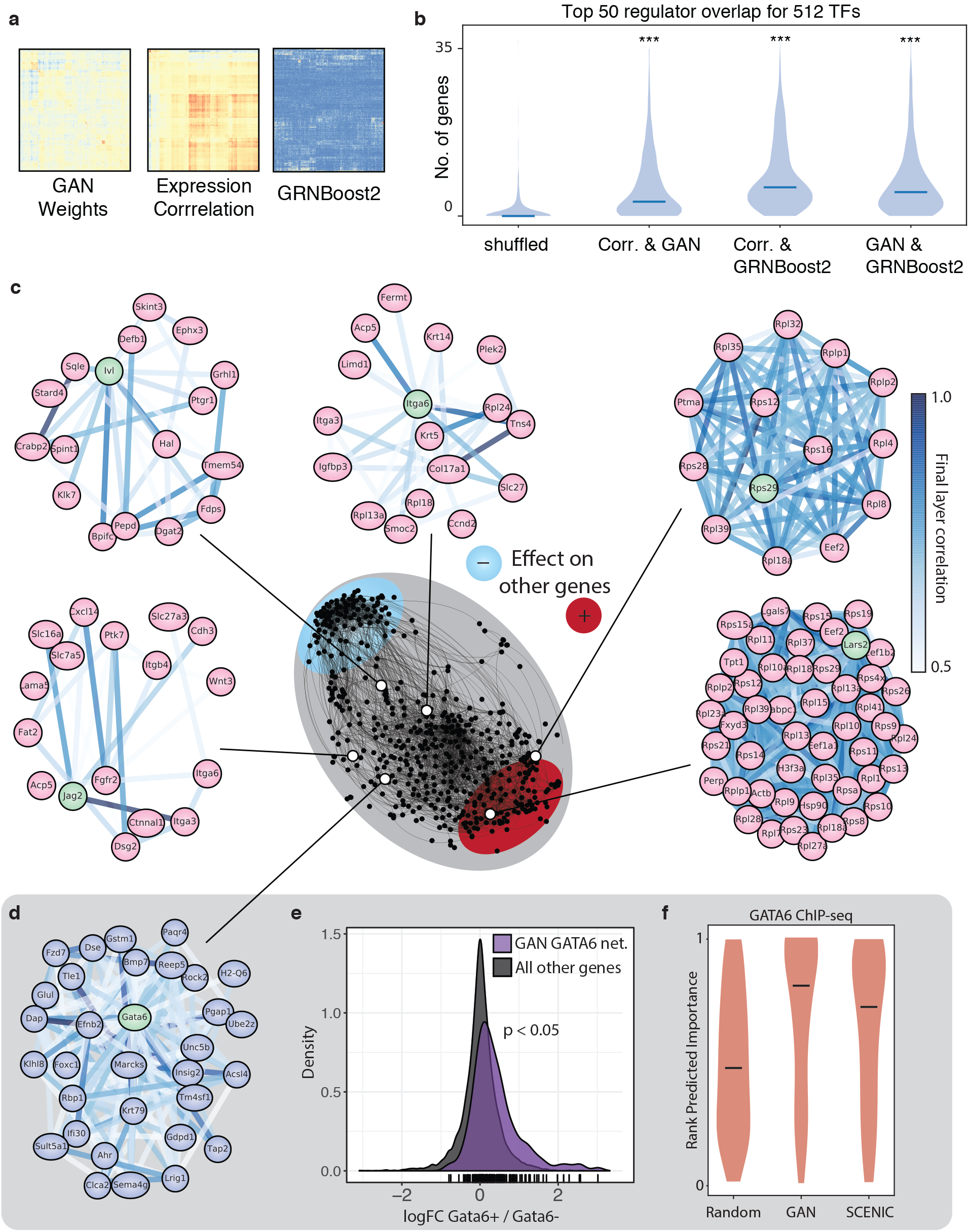
GAN-derived gene association networks. **a**) Gene-gene association scores derived from the generator neural network (left), gene coexpression correlation (middle) and GRNBoost2/SCENIC (right). **b**) Violin plots showing the distribution of overlap between the top 50 predicted target genes for 512 transcription factors. Shuffled shows the expected overlap distribution for randomly sampled genes. **c**) Network representation of the correlation between discriminator neural network final layer values for 6605 genes (centre panel); genes are depicted as nodes and correlations between them are shown as edges (threshold final layer correlation: 0.5). Network regions containing genes with overall positive and negative effect on other genes are highlighted in red and blue respectively. Surrounding local network are local gene association networks for Lars2, Itga6, Ivl, Rps29 and Jag2; edges are colored by generator final layer correlation. **d**) Local gene association network for Gata6 displayed as described above. **e**) Probability density plot of the log-fold change in gene expression between Gata6+ and Gata6^−^ cells for genes in the predicted GATA6 local association network (violet) and all other genes (grey). Network genes display greater expression changes between Gata6+ and Gata6^−^ cells (p-value = 1.3e-4). **f**) Violin plot showing the predicted rank importance of GATA6 targets as predicted by the GAN and SCENIC. Random shows a null distribution. GATA6 targets are defined by ChIP-seq from Wamaitha et al.

We visualised the correlation of the final layer weights for the 600 features between all genes using a force directed network (Figure 6C, middle network) and also examined the structure and clustering of the genes using a hierarchically clustered heatmap (Supplementary Figure S6). On a macroscopic scale genes are segregated by their overall positive or negative effect on gene expression. On a local scale the majority of genes cluster into small groups of between 10 and 50 closely associated genes. We used these local gene association networks to examine Lars2, a gene predicted from our analysis to be highly important for epidermal cell state. Lars2 is a mitochondrial leucyl-tRNA synthetase and our analysis shows it to be a member of an extremely closely regulated group of ribosomal and translation related genes. We also examined the local gene association networks for three known epidermal regulators examined in our previous sensitivity analysis Krt14, Itga6 and Ivl. Itga6 and Krt14 are both known basal IFE markers; the local network for these genes highlights the power of our machine learning approach to identify coregulated genes. Several known basal IFE regulators are present both local networks, such as Krt5, Krt14, Itga3, Cav2 and Col17a1. We observed a similar pattern of enriched associated genes for Ivl, an IFE differentiation marker which is co-associated with several other differentiation genes and Rps29 a marker of IFE-derived cells which is exclusively associated with ribosomal genes in our local networks. Furthermore, the local network for Notch ligand Jag2 contains several hair follicle bulge and basal IFE regulators, corresponding to its role in these epidermal locations^35^. Strikingly, Wnt3 is present in this local network indicative of Wnt-Notch signaling interplay^36,37^, which cannot be seen at the RNA level using conventional gene expression analysis. These co-regulated epidermal genes are robustly identified despite little correlation of expression in the scRNA-seq data.

Finally, in order to explore the predictive power of our neural network approach we derived the local gene association network for the GATA-binding factor 6 transcription factor, GATA6 (Figure 6D). We predicted that as a transcription factor, the GATA6 network would contain many genes which are directly regulated by GATA6. Hence, we hypothesised that genes in the network with a positive final layer correlation should be upregulated in GATA6+ cells. Using a previous dataset from the lab, we contrasted bulk gene expression of GATA6+ and GATA6^−^ cells derived from the junctional zone and sebaceous duct of the epidermis (Supplementary Figure S8)^38^. Figure 6E shows the log fold-change distribution of genes between the GATA6+/GATA6^−^ subpopulations. Genes derived from our GATA6 gene regulatory network show significantly higher expression in the GATA6+ cells when compared to all other genes (p < 0.05, Kolmogorov-Smirnov test). Finally, we sought to compare the ability of the GAN algorithm and SCENIC to identify Gata6 targets. We used Gata6 ChIP-seq data from Donati and Wamaitha to define direct targets of Gata6 and for each target extracted the rank association score derived from the GAN and SCENIC gene regulatory networks (Figure 6F)^39^. GAN and SCENIC both successfully predict high association scores and are statistically significant relative to a random control sample of genes. Furthermore, the GAN algorithm predicts a higher median score for these targets relative to SCENIC. This comparison confirms that both methods are similarly able to predict Gata6 targets, however, for the GAN these associations are obtained as a byproduct of a joint generative and discriminative model. Taken together these results suggest GANs are capable of elucidating complex gene interrelationships beyond the limits of linear correlation analyses.

### 2.6 Discriminator network + t-SNE allows better dimensionality reduction

Finally, we hypothesised that the discriminator network learned biologically relevant features of scRNA-seq data, discarding uninformative genes in order to discriminate successfully between the generated and real samples. The discriminator neural network consists of a single hidden layer whose output is transformed to a discriminator or “critic” output value It transforms 6605 gene expression values into 200 learned internal features, effectively a reduced dimensional representation. Hence, we sought to understand whether the discriminator hidden layer output contained learned features that could be used to improve on existing dimensionality reduction. We performed t-SNE on the discriminator hidden layer output to visualise these learned features (Figure 7) and compared this with two other approaches.

**Figure 7.**
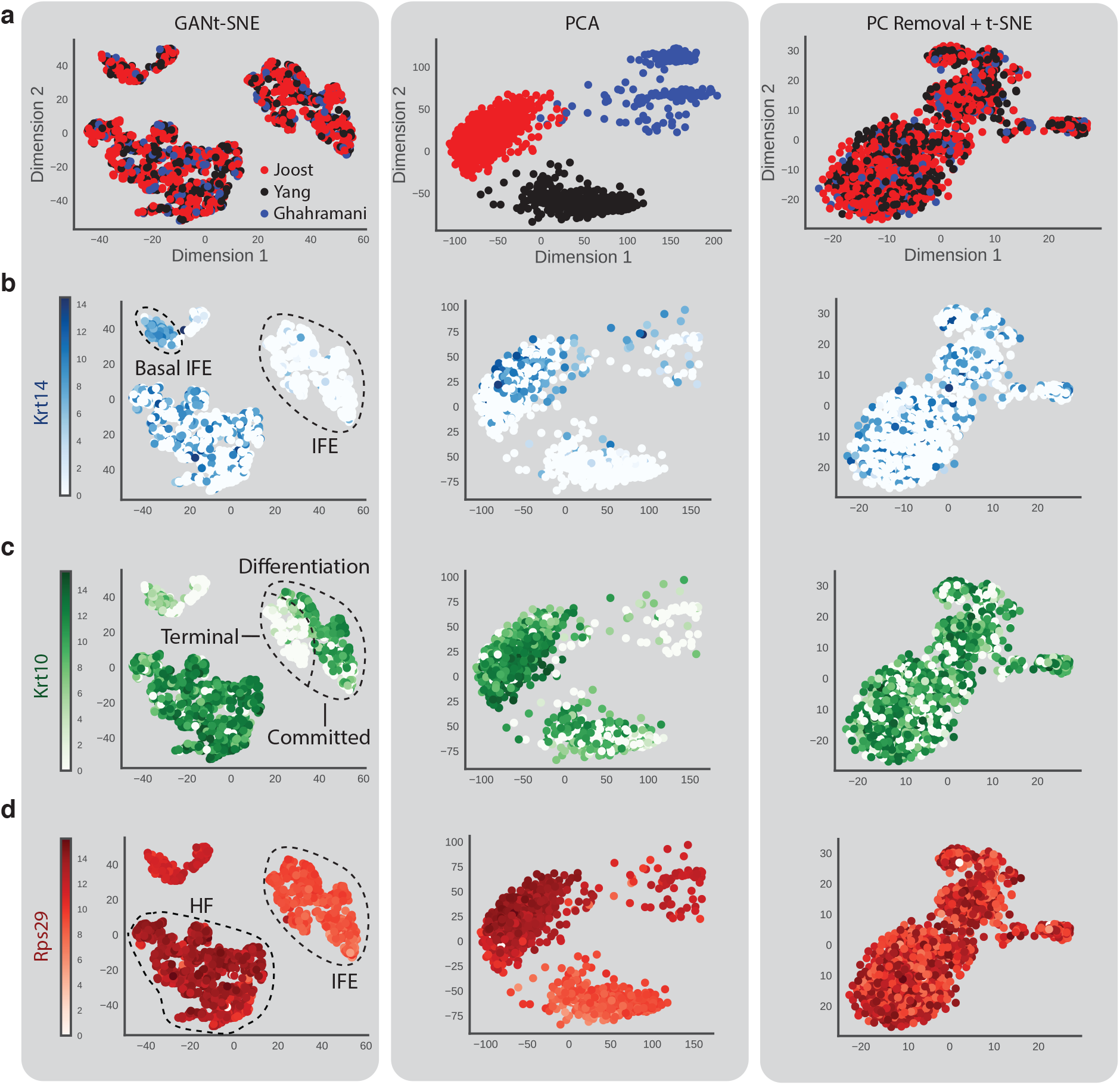
GANt-SNE clusters biologically similar cells. **a**) GANt-SNE visualises 2263 real cells by performing t-SNE on 200 features captured by the discriminator neural network’s hidden layer. This is compared with principal component analysis (PCA) and t-SNE after removal of linear batch effects using PCA (PC removal + t-SNE). PCA is heavily affected by batch effect and clusters the three datasets separately. **b**) Keratin-14 expression overlaid onto projections. Krt-14 is a marker for basal interfollicular epidermis (IFE). GANt-SNE distinguishes between basal IFE and non-basal IFE cells, whereas PCA and PC removal + t-SNE fail to distinguish between these cell types. **c**) Keratin-10 is a marker for differentiation interfollicular epidermis (IFE). GANt-SNE successfully clusters separates supra-basal and terminally differentiated IFEs. **d**) Ribosomal Protein S29 is known to be expressed highly in interfollicular epidermis (IFE) and at medium levels in hair follicles (HF). GAN-tSNE successfully distinguishes between these two major cell types.

First we performed a principal component analysis (PCA) on the combined epidermal training data (Figure 7A, middle column); the main source of linearly separable variance in the first two components is the batch effect of dataset origin and laboratory and the approach generally fails to identify biologically meaningful clusters even at higher principal components. We removed this technical source of variation by subtracting the first two principal components and performing a t-SNE on the resulting dataset. The resulting t-SNE (Figure 7A, right column) no longer clusters cells by dataset origin, so removing variation caused by different experimental protocols, at the cost of concurrently removing informative biological variation in gene expression.

Two major sources of variation in epidermal cells are their differentiation status and spatial position within the epidermis: a useful dimensionality reduction approach should capture at least one of these features, ideally both. To investigate this we overlaid expression of Krt14, a marker of undifferentiated keratinocytes, Krt10 a marker of commitment to differentiation and Rps29, a ubiquitously expressed ribosomal protein upregulated in the IFE in comparison to hair follicles.

Biologically, we expect cells with similar expression levels of these marker genes to cluster. PCA and the combined approach of PC removal + t-SNE perform poorly in this regard and expression of all three markers fails to distinguish cells across the dimensionally reduced space (Figure 7 middle and right column). In contrast, our GANt-SNE approach clusters Krt14^high^ cells deriving from the basal layer of the IFE (Figure 7B, Basal IFE) which are separated from the remaining IFE cells. A third cluster of hair follicle cells shows sporadic expression of Krt14 as this is not a functional marker for these cells. Our dimensionality reduction separates differentiated IFE cells highly expressing Krt10 from terminally differentiated and keratinized Krt10^low^ cells (Figure 7C). Furthermore, GANt-SNE correctly separates IFE-derived cells from hair follicle cells as seen by Rps29 expression (Figure 7D). These biological distinctions are not identified by the two alternative methods of dimensionality reduction.

These results lend credence to our hypothesis that the discriminator learns biologically relevant features of the data. Using the GANt-SNE approach we successfully separate cells by differentiation status and spatial position (IFE vs. hair follicle and differentiating vs. undifferentiated). This is achieved without a priori knowledge of technical variation and batch effects by training on the commonalities between training data. In contrast the PCA and t-SNE approaches fail to separate cells into these biologically meaningful clusters, as they crudely extract the strongest sources of variation - often a non-linear mixture of biological and technical variation.

## 3. Discussion

We have addressed a major goal of single cell gene expression analysis; the desire to obtain functional gene relationships and a predictive model of transcriptional state in single cells. This serves as a framework for understanding cell fate transitions, disease-causing perturbations and state-determining genes across tissue types.

Here we demonstrate the effectiveness of a new deep learning approach in integrating multiple gene expression datasets, despite no prior adjustment for technical and batch-to-batch variation. To the best of our knowledge, this is the first time GANs have been applied to genomic data. Our data indicate that GANs are able to accurately simulate gene expression relationships in single cells. Simulation of gene expression alone will lead to a better understanding of single cell variability and improve our ability to predict the effects of cellular perturbations.

In comparison to other scRNA-seq analysis methods, GANs uniquely combine multiple desirable features such as data integration, prediction of gene expression dynamics, gene-gene association networks, and dimensionality reduction. Alongside these features, a trained GAN provides a generative model - applications of which are yet to be fully explored. Our approach of combining data from cells in different environments with diverse technical and biological noise allows robust inference of gene associations intrinsic to epidermal cells rather than experimental conditions. We focused analysis and exploration of the capabilities of GANs on skin due to the wealth of knowledge available regarding subpopulations and cell state. However, our generative model can be extended to integrate cells from multiple tissues and we anticipate this will advance our understanding of gene expression relationships across tissues.

As further scRNA-seq data becomes available, particularly through large scale projects such as the Human Cell Atlas^40^ we envisage that GANs will be a viable strategy to analyse all human cell types en masse. In our study we have shown that a relatively small dataset of under 2000 epidermal cells is sufficient to uncover state-determining genes and gene expression networks in the epidermis. GANs applied to a dataset covering multiple tissues could be used to resolve organism-wide gene expression relationships and to predict previously unseen cell state transitions.

## 4. Methods

### Datasets and data preparation

All datasets analysed in this study are publicly available under the GEO accessions GSE90848 (Yang, mouse hair follicle subpopulations), GSE67602 (Joost, mouse epidermal cells), GSE99989 (Ghahramani, *in vitro* mouse keratinocytes), GSE96772 (Buenrostro, human blood cells), at the Short Read Archive under accession number SRP073767 (Zheng, human blood monocytes) or at ArrayExpress under accession E-MTAB-5466 (Sagner, mouse neural progenitors). We combined the three epidermal datasets (GSE90848, GSE67602, GSE99989) using transcripts per million (TPM) normalisation and removed labelled non-epithelial cells from the Joost dataset. Data was filtered for: cells with more than 1000 genes detected log_2_(TPM+1) > 1 and genes expressed in more than 500 cells at log_2_(TPM+1) > 1. All neural network input data was log_2_(TPM+1) and the combined dataset is available as a CSV matrix at https://github.com/luslab/scRNAseq-WGAN-GP/tree/master/data.

### Tools and code availability

Data preparation was performed using R. Gene ontology enrichment was evaluated using enrichR^41^. All other analysis and implementation of the generative adversarial networks was performed using Python (numpy, sklearn and networkx). We used Google’s Tensorflow^42^ deep learning framework to implement, train and monitor our neural networks. We have provided a Jupyter Notebook containing our implementation of the generative adversarial network available at https://github.com/luslab/scRNAseq-WGAN-GP.

### Generative adversarial network algorithm

We adapted our generative adversarial network algorithm from four previous works^9,10,13,24^. Both generator and discriminators are fully connected neural networks with one hidden layer. We used a Leaky Rectified Linear Unit (LReLU) as the activation function for both networks using a coefficient of 0.2.

The generator input consists of a 100-dimensional latent variable (z) and a hidden layer size of 600 fully connected units. In order to reduce training time and data required for the generator network to simulate the Poisson distributed nature of scRNA-seq count data we trained the generator network using an additive Poisson and Gaussian distributed latent variable where *z* = *N*(*μ* = 0, *σ* = 0.1 * *max*) + *P*(*λ* = 1). Generator output is a 6605 dimensional vector representing the 6605 genes in the training cohort.

The discriminator network input is a 6605 dimensional vector (gene expression profile) and has a hidden layer size of 200 fully connected units. The final layer of the discriminator does not apply an activation function, in line with other Wasserstein GANs using a gradient penalty loss function.

We used backpropagation and RMSProp to train the two neural networks (learning rate of 5e-5) and a batch size of 32 cells. Additionally, data were augmented during training by randomly permuting the expression of 10 genes expressed at log_2_(TPM+1) < 3. We used several loss functions during this study before finalising on using the Wasserstein-GAN (WGAN) with gradient penalty loss function. WGAN replaces the original GAN where data distributions of generated and real data are compared using the Jensen-Shannon divergence for the loss function^9^. In WGANs the loss function is calculated using the Wasserstein distance which gives improved training stability^13^. Gulrajani and colleagues further improved on this variant of GANs by eliminating weight-clipping in favor of a gradient penalty in the form of the gradient between pairs of real and generated samples^24^.

The two loss functions to minimise are:

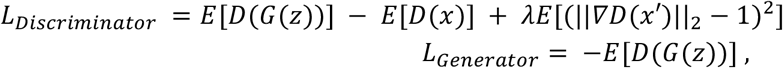

where *G* is the generator network and Dis the discriminator network such that 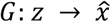, *z*is the generator latent variable, 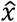 is a generated gene expression profile, *x* is a real gene expression profile and *x*′ is uniformly sampled between pairs of generated and real gene expression profiles.

Convergence of the loss function is not sufficient to evaluate completed training of GANs. For each timepoint we evaluated the diversity of cells produced by the generator by simulating 500 cells and calculating the median distance (Pearson), i.e. median cell-cell distance.

For a list of GAN parameters used see Supplementary Table 2.

### Dimensionality reduction and clustering

We evaluated clustering and structure of real and generated single cell RNA-seq profiles in three ways 1) PCA (principal component analysis), 2) t-distributed stochastic neighbour embedding (t-SNE) on data with the top two principal components removed. 3) t-SNE on the hidden layer output values of the discriminator network. For all t-SNEs we used a default perplexity parameter of 30.

Simulated time-series gene expression profiles were clustered by k-means clustering with 20 expected clusters. Clustered dynamic gene expression profiles were subsequently visualised and two pairs of clusters were manually merged where they were deemed to be visually similar.

### Latent space mapping and interpolation

Terminally differentiated cells and basal IFE cells were sampled from the combined dataset on using labels from the original studies. In order to generate corresponding latent space variable *Z_basal_* and *z_differentiated_* for cells *x_basal_* and *x_differentiated_* we randomly generated cell expression profiles until the correlation between x and the generated cell G(z)reached a threshold of 0.7. Figure S1B shows that a correlation of 0.7 between two cells is rare, hence, we assume that this threshold is sufficient to generate a gene expression profile representative of the original cell state. We next calculated the difference between the basal and differentiated cell in latent space i.e. *S* = *z_basal_ - z_differentiated_*. This latent space representation of differentiation was used to simulate differentiation on a further unseen cell where we obtained *z_unseen_* and calculated a predicted terminally differentiated latent space point *Z_unseen differentiated_ z_unseen_* + *δ*. To obtain simulated gene expression time series data we interpolated 1000 points between *z_unseen_* and *z_unseen differentiated_* and generated 1000 intermediate gene expression profiles from these.

For Monocle analysis, we used the subset of cells identified as from the IFE in Joost et al. Monocle was run using default settings to obtain dimensionality reduction, pseudo-ordering and subpopulation clusters.

### Discriminator network sensitivity analysis

Contribution of genes to output and sensitivity of the discriminator network was performed by taking unseen cells and varying the output of each gene between the minimum and maximum observed across all cells. For important genes this results in a change of the discriminator output *D*(*x_p_*) where *x_p_* is a perturbed gene expression profile. We performed 100 linearly interpolated perturbations. Sensitivity curves were calculated by taking the mean normalised *D*(*x_p_*) at each expression level as unseen cells have can differ in their baseline or unperturbed discriminator score *D*(*x*). Max sensitivity gradient (see Figure 5B) was calculated by taking the absolute value of maximum gradient of these curves.

### Local gene association networks

Gene co-regulatory relationships were inferred by analysing the weights of the generator neural network final layer. The Pearson correlation matrix of the final layer was used as a network adjacency matrix. To construct association networks for a gene we selected all local genes with a final layer correlation > 0.5 and retained any connections within the local network also with final layer correlation > 0.5.

## Author Contributions

AG performed analyses and wrote the manuscript. NML and FMW supervised the project and co-wrote the paper.

## Acknowledgements

We are grateful to Victor Augusti Negri, Aylin Cakiroglu and Dean Plumbley for helpful advice and discussions. This work was supported by the Francis Crick Institute which receives its core funding from Cancer Research UK (FC010110), the UK Medical Research Council (FC010110), and the Wellcome Trust (FC010110), Wellcome Trust funding to NML (103760/Z/14/Z), and MRC Medical Bioinformatics Award eMedLab to NML (MR/L016311/1).

---

**Supplementary Figure S1.**
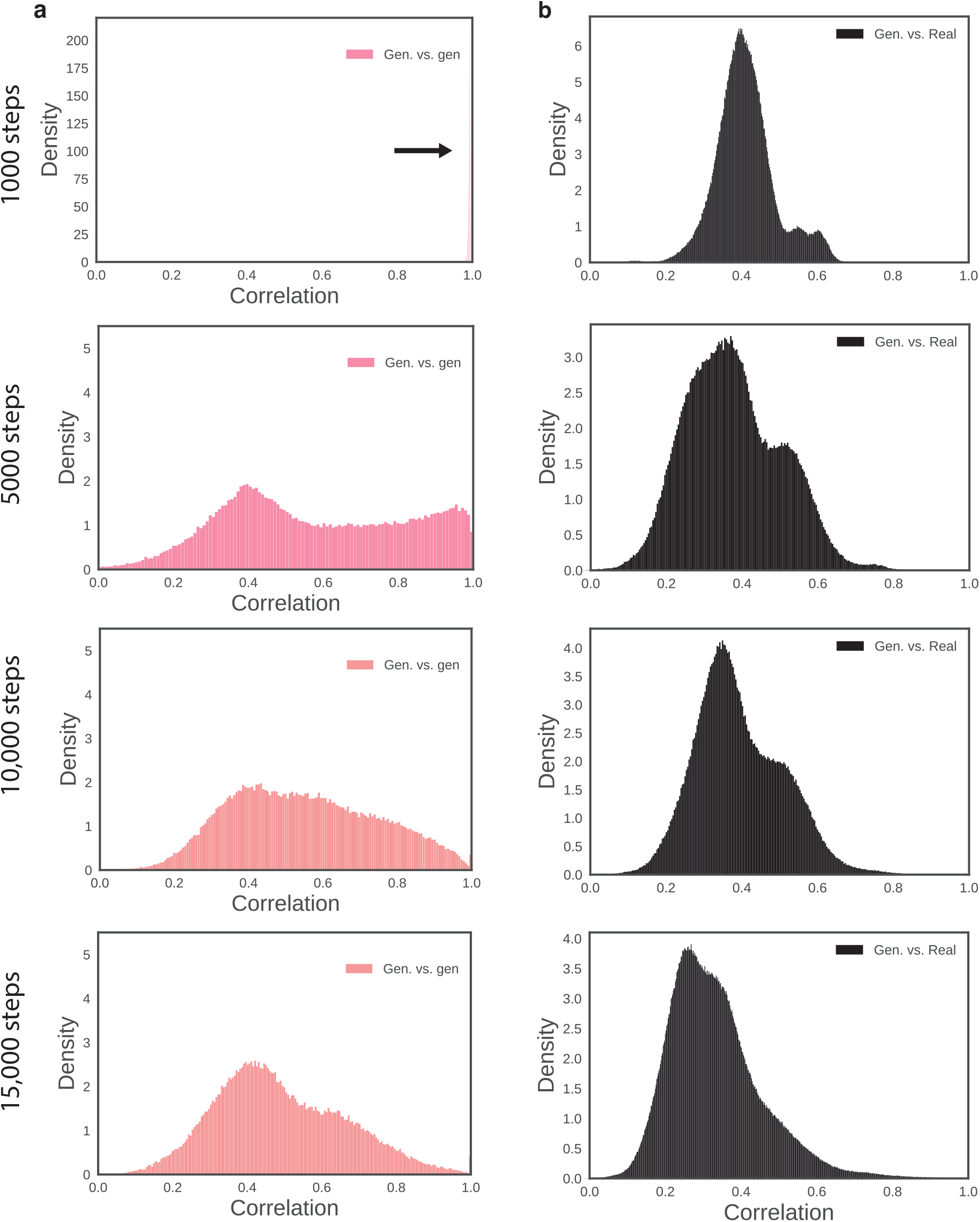
GAN training improves expression output diversity. **a**) Distribution of pairwise correlation between pairs of real or generated cells after 1000, 5000, 10,000 and 15,000 training steps. **b**) Distribution of pairwise correlation between pairs of real and generated cells after 1000, 5000, 10,000 and 15,000 training steps.

**Supplementary Figure S2.**
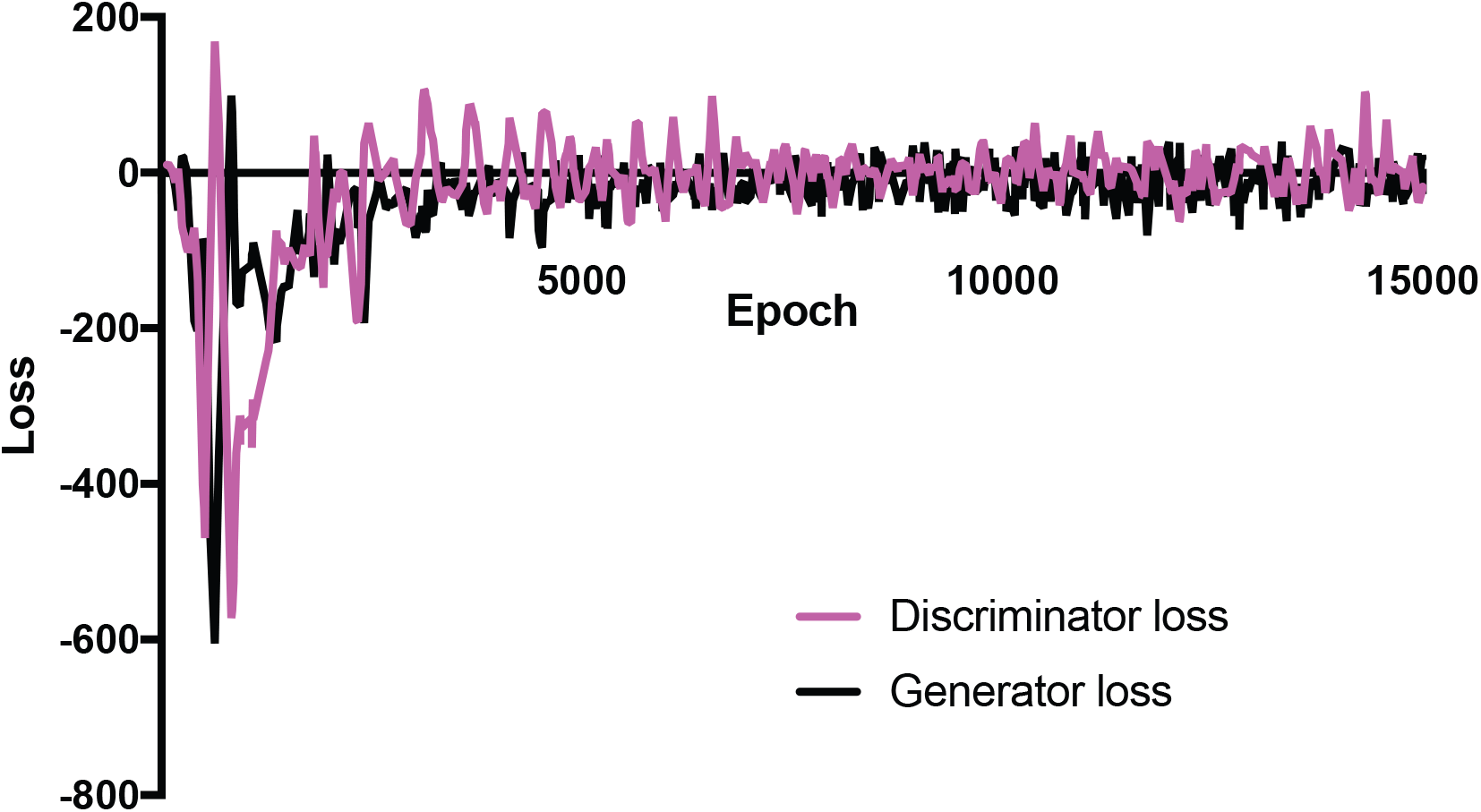
Generator and discriminator network loss curves. Pink line shows discriminator network loss curve, black line shows generator network loss curve. Unlike most neural networks, convergence of loss for GANs is not sufficient to indicate that training is complete.

**Supplementary Figure S3.**
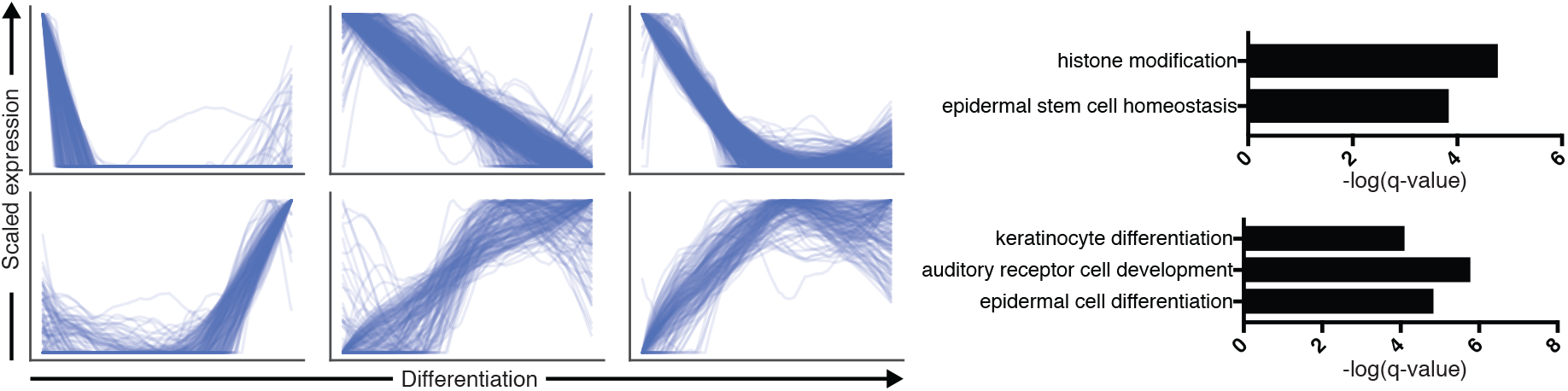
Gene ontology enrichment for predicted differentiation genes. Gene ontology enrichment for genes that generally decrease (upper) or increase (lower) over differentiation as predicted by latent space interpolation.

**Supplementary Figure S4.**
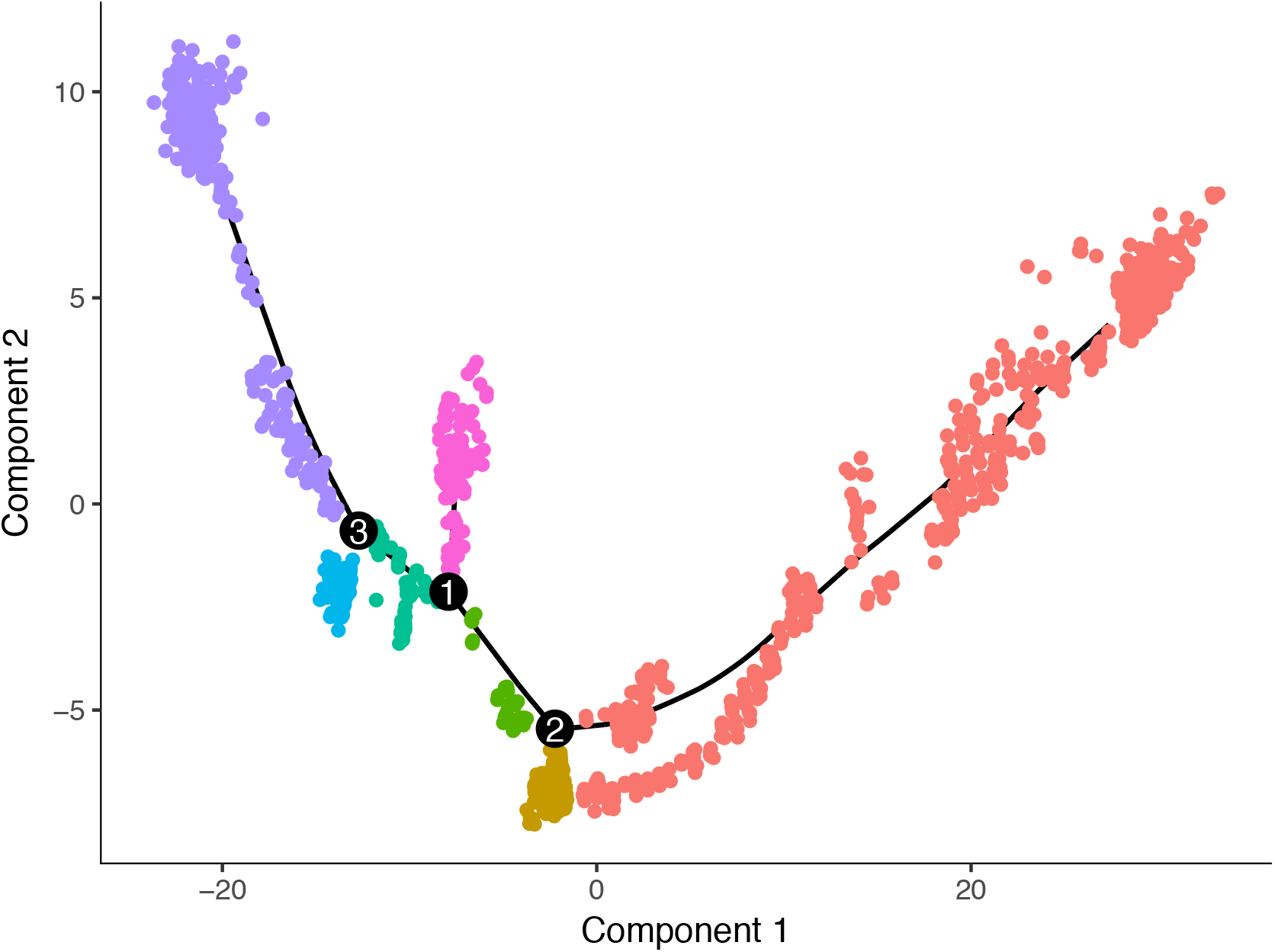
Dimensionality reduction and pseudo-ordering of IFE cells from Joost et al. using Monocle to reconstruct differentiation gene expression dynamics in the IFE.

**Supplementary Figure S5.**
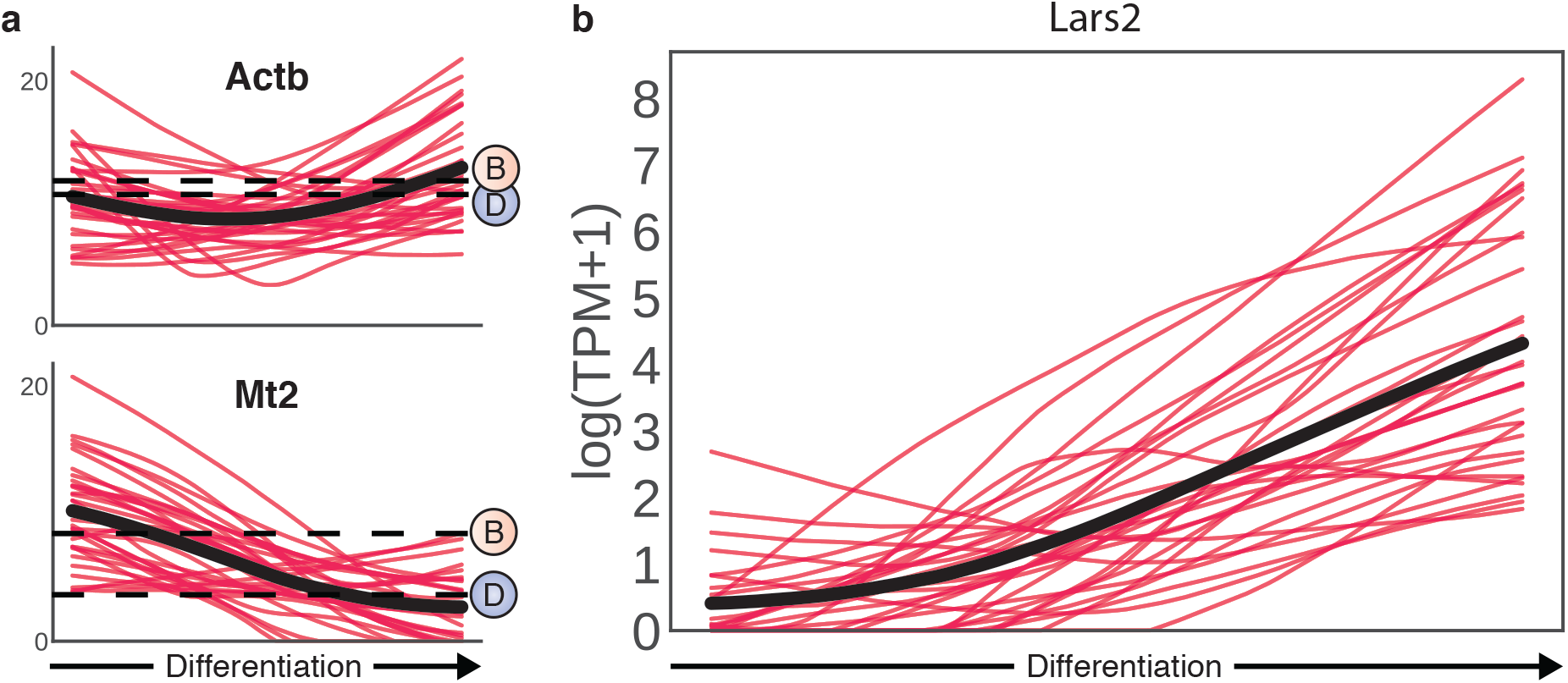
GAN-simulated gene expression simulated over differentiation. Lars2, Actb and Mt2 expression over differentiation for 30 generated single cells. Red lines are 1000 simulated differentiation timepoints. Black line represents mean expression over all cells.

**Supplementary Figure S6.**
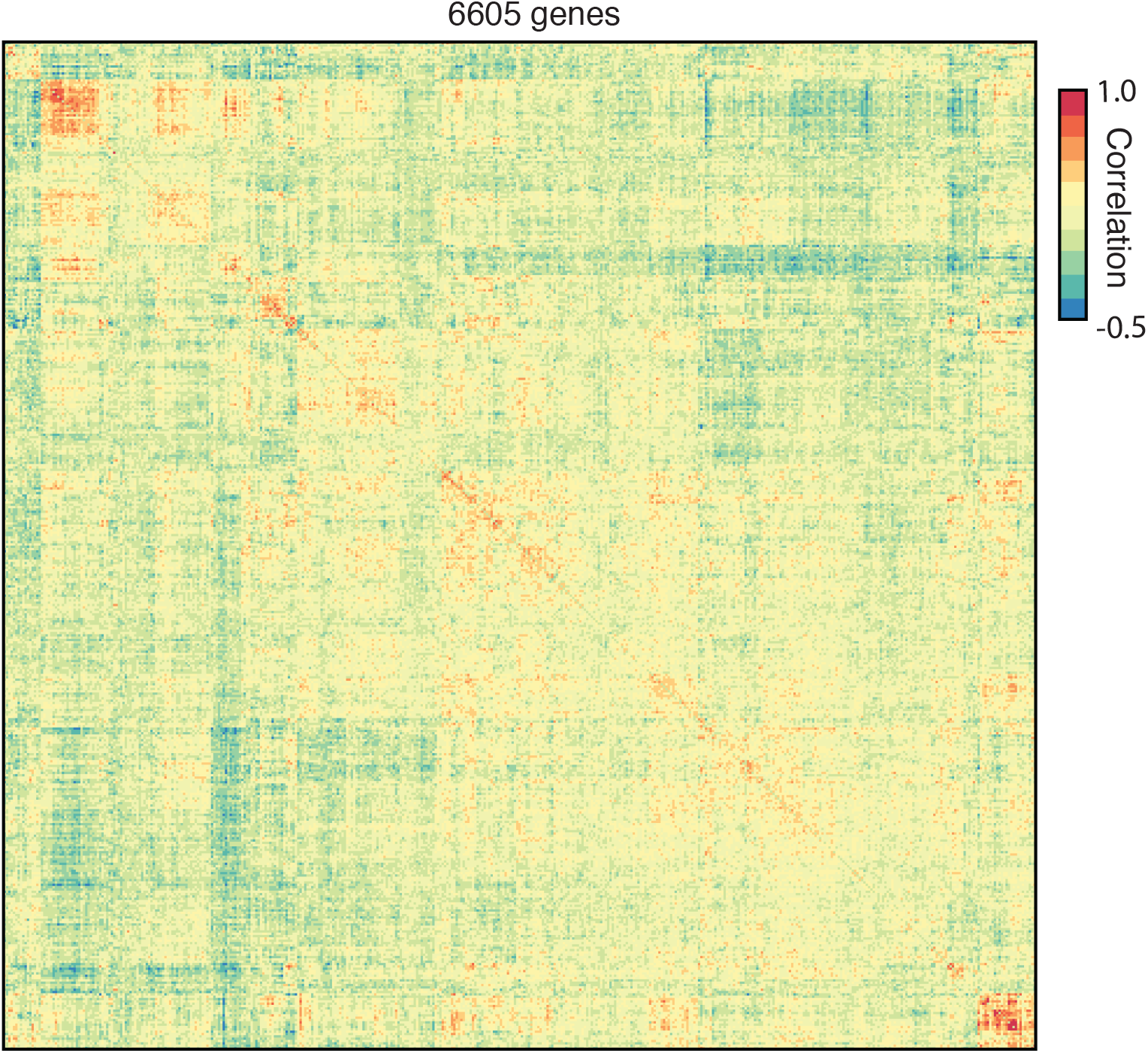
Correlation matrix of the generator neural network final layer weights visualised using a heatmap (Pearson correlation).

**Supplementary Figure S7.**
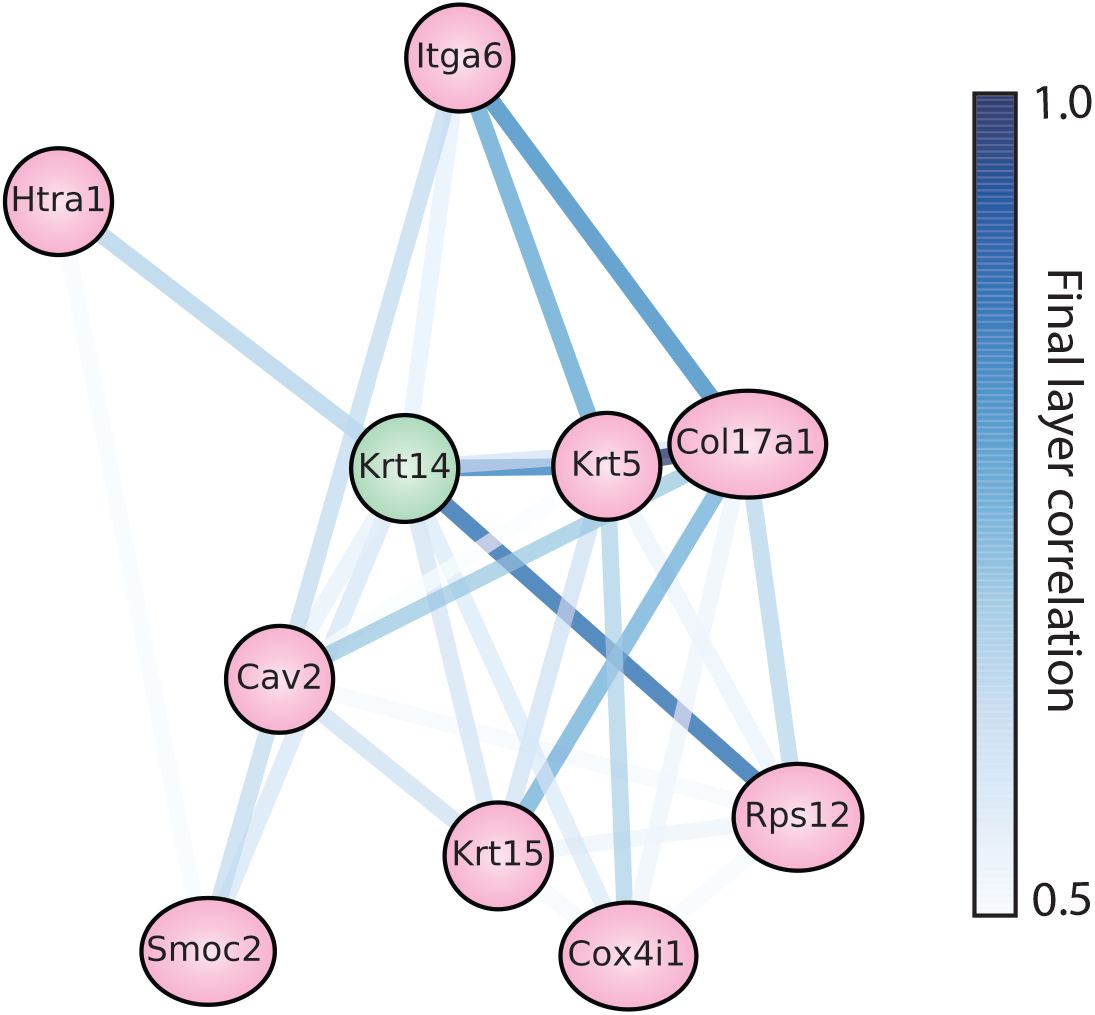
Local gene association networks for Krt14 visualised using a Fruchterman-Reingold force-directed network. Edges are colored by generator final layer correlation.

**Supplementary Figure S7.**
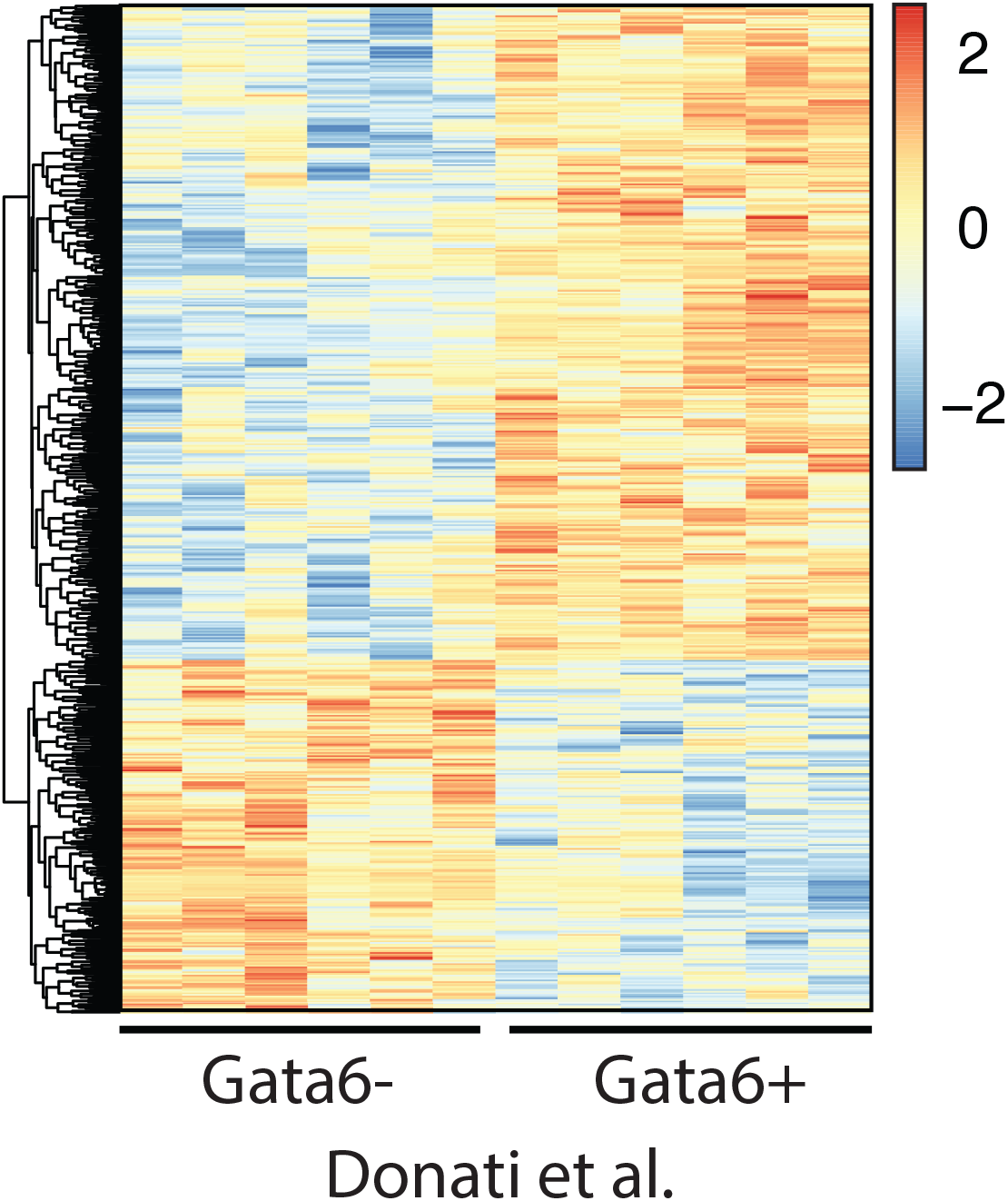
Heatmap showing differentially expressed gene expression z-scores for Gata6+ and Gata6^−^ junctional zone and sebaceous duct cells from Donati et al.

## Supplementary Tables

Supplementary Table 1. Genes identified by sensitivity analysis.

List of genes ranked by maximum discriminator network sensitivity gradient.

Related to Figure 5B

Supplementary Table 2. List of GAN neural network parameters.

